# Trauma-Exposed Adolescents Show Reduced Cortical Glutamate Modulation during Inhibitory Control with Negative Emotional Stimuli

**DOI:** 10.64898/2026.07.03.735903

**Authors:** John McClellan France, Dalal Khatib, Shaurel A. Valbrun, Sattvik Basarkod, William M. Davie, Manessa Riser, Vaibhav A. Diwadkar, Noa Ofen, Hilary A. Marusak, Ana M. Daugherty, Tanja Jovanovic, Jeffrey A. Stanley

## Abstract

**Background:** Childhood trauma exposure (TE) may heighten negative emotional responses, overwhelm cognitive control, and increase risk for anxiety disorders. Cognitive control is facilitated by glutamatergic (Glu) excitatory neurotransmission within the dorsal anterior cingulate cortex (dACC). Dynamic changes in dACC Glu levels were investigated using ¹H functional magnetic resonance spectroscopy (^1^H fMRS) to assess the impact of negative emotional processing on neural mechanisms supporting cognitive control in TE-youth.

**Methods:** Fifty adolescents were categorized into two TE-Groups: Higher (Mtrauma*=6*±1events) and Lower (Mtrauma*=3*±1events). ^1^H fMRS from the dACC was acquired during an inhibitory motor control task requiring tapping responses to stimuli under two Response Modes, “NonSelective” (100% response) and “Selective” (80% response, 20% inhibition), executed with two Stimuli Conditions, “Squares” (no emotion) and “Faces” (emotion). Glu modulation (relative to basal levels) was tested across TE-Group, Stimuli Condition, and their interaction. Within each Stimuli Condition, Glu modulation was tested across Response Modes by TE-Group.

**Results:** We observed a 2-way interaction of TE-Group x Stimuli Condition (χ^2^=4.66, *p*=0.031). Post-hoc tests revealed significantly lower Glu modulation in Higher TE vs Lower TE (*p*=.023) during Faces but not Squares. This Glu modulation did not differ across Response Modes. Within the Higher TE-Group, Glu was significantly reduced during Faces compared to Squares (p<.001). Basal dACC Glu levels did not differ between groups.

**Conclusions:** TE-Group differences in adolescent dACC Glu modulation were observed during cognitive control performed with emotional, but not non-emotional, stimuli, highlighting the value of ^1^H fMRS for detecting trauma-related differences in task-related excitatory neurochemical dynamics.

## Introduction

Childhood trauma exposure (TE), including exposure to actual or threatened death, serious injury, or sexual violence, is highly prevalent in the United States^1,2^ and can greatly increase the risk for developing anxiety disorders ^3^. Notably, anxiety disorders frequently emerge during adolescence^4^, implicating adolescent neurodevelopment in the association between TE and anxiety. Developmental neurobiological theories posit that TE can heighten emotional lability, leading to exaggerated emotional responses to affective stimuli^5–7^. In turn, heightened emotional responses may overwhelm cognitive control processes (i.e. bottom-up emotion interference of top-down cognitive control). This disruption may affect emotion regulation and healthy goal-driven behaviors, and thereby contribute to anxiety symptomology^8–10^. Consistent with these theories, TE in adolescents has been associated with exaggerated emotional responses to potential threats and stressors^11–13^, elevated and prolonged activation in emotion-related brain regions^14–17^, and altered activity in prefrontal cortex during cognitive control tasks involving emotional stimuli^18–20^. Accordingly, TE adolescents may be particularly vulnerable to emotion interference of cognitive control, underscoring the need to identify the neurobiological mechanisms underlying this vulnerability and their contribution to anxiety symptoms.

Inhibitory control is defined as a cognitive control process involving suppression of prepotent behaviors that are incompatible with internal goals^21^, and inhibitory control deficits have been associated with fear and anxiety-related pathology^22,23^. While present during early development, inhibitory control greatly improves during adolescence before reaching adult-levels^24–26^. Emotion interference of inhibitory control can be investigated when performance is challenged by salient emotional distractors, such as emotional face stimuli. Emotional go/no-go tasks embed emotional face stimuli directly into the foreground of a standard go/no-go paradigm^27^. In such tasks, negative emotional faces, unlike positive faces, selectively impact behavioral accuracy in trauma-exposed youth compared to non-trauma exposed youth^8^, providing evidence for negative emotion interference of inhibitory control.

Converging research suggests that inhibitory control is facilitated in part by top-down cognitive control regions of the cortex, including the dACC, which communicates with the basal ganglia to execute and modify planned motor behaviors^21,28^. The dACC is particularly well-positioned to convey top-down information flow because it shares anatomical projections from the lateral prefrontal cortex and to the motor cortex^29^. Functional MRI studies have provided extensive evidence that implicates the dACC in inhibitory control in both adolescents and adults^30–32^, with electrical brain stimulation and brain lesions providing further support for the region’s causal role in the initiation and execution of motor movement^33,34^. Moreover, changes in dACC activation mediate longitudinal improvements in inhibitory control during adolescence^24^. Importantly, TE has been linked to developmental changes in dACC structure and function^35^. Together, these findings position the dACC as an apex region for top-down inhibitory control and suggest that TE-related differences in this region may contribute to emotion interference of inhibitory control.

Despite evidence for the neurochemical basis of brain function^36–38^, the neurochemical mechanisms underlying dACC activation remain poorly characterized. The activity of the dACC is mediated by the interaction between excitatory and inhibitory neurons, principally reliant on glutamate (Glu) and gamma-aminobutyric acid (GABA) neurotransmission, respectively. The dynamic interplay of Glu and GABA signaling is tightly coupled^39–41^ and supports coordinated neural activity that is essential for cognitive control^42–44^. Importantly, traditional methods to study brain function like functional MRI measure hemodynamic responses which are downstream of Glu and GABA signaling^45^. Thus, the neurochemical mechanisms that underpin inhibitory control, and the impact of TE on these mechanisms, remain largely unknown.

^1^H functional magnetic resonance spectroscopy (^1^H fMRS) measures dynamic changes in steady-state Glu (and/or GABA) levels across task conditions (i.e., modulation) *in vivo*^36,46,47^ and therefore is a useful tool to investigate the neurochemical processes underlying inhibitory control. ^1^H fMRS has been used in healthy adults performing a Stroop task which led to an approximate 2.5% increase in dACC Glu level (averaged across the complete task) relative to a preceding non-task comparison condition^48^. A follow-up investigation using the same Stroop task demonstrated differences in task-related Glu modulation in adults with schizophrenia and depression compared to healthy controls^49^. These findings suggest that in healthy adults, inhibitory control is associated with a positive Glu modulation, which can be impaired in psychopathologies that are associated with inhibitory control dysfunction. It remains unknown if emotion interference disrupts dACC Glu modulation during inhibitory control in adolescents with TE, a high-risk developmental period for TE and TE-related psychopathology.

To address this gap, the current study used ^1^H fMRS to measure dynamic changes in dACC Glu levels using a 2×2 block design, visually guided inhibitory motor control task in adolescents with TE. The task included two Response Modes: 1) NonSelective, which involved only motor control without inhibition (i.e., all-go task blocks) and 2) Selective, which involved both motor control and inhibition (i.e., go/no-go task blocks). Both Response Modes were executed across two Stimuli Conditions; 1) Squares, or non-emotional stimuli, and 2) Faces, or negative and neutral emotion stimuli, to assess the emotion interference of cognitive control. We hypothesized that youth who endorsed a greater degree of trauma exposure, compared to youth who endorsed a lesser degree, would demonstrate lower Glu modulation during the Faces Stimuli Condition but not Squares, which would be consistent with an emotion interference effect. We also hypothesized that any trauma-related differences in Glu modulation during Faces would be amplified during Selective responding compared to NonSelective responding, which would be consistent with emotion interference of inhibitory control, specifically. Finally, we hypothesized that Glu modulation during the Faces condition would be negatively associated with anxiety symptom severity.

## Methods

### Participant Demographics

A sample of 53 participants between the ages of 10 and 15 years old (27 females and 26 males, m*ean age* ± SD: 13.3 ± 1.5 yrs) were recruited from Detroit, MI, through flyer distributions at local libraries and community centers. Individuals who endorsed diagnosis of a current or past DSM-5 Axis I psychiatric disorder including an anxiety disorder, and/or currently receiving psychotropic medication (e.g., an antidepressant/mood stabilizer/psychostimulant) or behavioral therapy were excluded to ensure a non-clinical sample. Additionally, individuals were excluded if they endorsed red/green color blindness, a neurological disorder, a developmental disorder, or a MR contraindication. All study procedures were approved by the Wayne State University Institutional review board (IRB000093684). Both the participant and their primary caregiver (i.e., parent or legal guardian) provided written informed consent and/or verbal assent. All study procedures were completed at a single visit.

### Trauma Exposure and Related Psychopathology Measures

All measures were administered verbally in English in an interview format by trained research staff to all youth participants without the presence of caregivers. Our prior work indicates that youth-reported trauma and symptoms exhibit stronger associations with neurobiological correlates than caregiver-reported measures of their child^50^.

#### Trauma Exposure

The Traumatic Events Screening Inventory (TESI) was used to assess participants’ self-reported exposure to potentially traumatic events^51,52^. The TESI is a 17-item questionnaire and has been validated against the Adverse Childhood Experiences (ACEs) measure^53^. Participants indicated whether they had experienced or witnessed various interpersonal (e.g., violence, abuse) or non-interpersonal (e.g., natural disaster) traumatic events. Trauma exposure (TE) was calculated as the total number of endorsed events, consistent with prior work^14^. A median split based on TE was applied to the total sample to establish the Higher and Lower TE-Groups (*Mdn*=4). Prior epidemiological studies conducted with nationally representative samples of children and adolescents suggest between a 22.9 – 49.5% prevalence rate for experiencing more than 1 traumatic event^54–56^, and that experiencing 4 or more adverse or traumatic events is associated with the greatest risk for psychopathology^54,57,58^.

#### Anxiety Symptoms

Child’s self-reported frequency of anxiety symptoms over the past 3 months were assessed using the Screen for Child Anxiety Related Emotional Disorders (SCARED)^59^, a 41-item scaled utilizing a 3 point frequency scale (0=Not True or hardly ever true,1=Somewhat true or sometimes true,2=Very true or often true). Anxiety severity was computed as a total of the frequency scores across all items, which has been shown to demonstrate good internal consistency and discriminant validity^59^.

#### Visually Guided Motor Inhibitory Control Task

This 2×2 block design included 2 Response Modes (i.e., NonSelective and Selective) executed across 2 stimuli type conditions (i.e., Squares and Faces), yielding 4 distinct task conditions. Conditions were executed as separate ¹H fMRS task runs (3:12 min/run) (**Figure 1a**). Task runs were counterbalanced across Response Modes, with participants completing either the NonSelective or Selective mode first. Each task run consisted of four 32-second working blocks, interspersed by 16-second rest epochs (fixation crosshair). The Squares stimuli included green or red Squares, and the Faces stimuli included negative or neutral faces selected from the NimStim set^60^. All stimuli were sized at 3.75” x 5.00” and were presented individually and in sequence for 300ms. Stimulus onset asynchrony was jittered across measurements (mean = 1231ms, range of 800ms – 1800ms, minimum response time 800ms), which is sufficient to induce a prepotent response while still allowing for accurate emotion discrimination^27^. Throughout the task, participants were instructed to respond as quickly and accurately as possible by tapping their right forefinger on an MR compatible tapping pad (Lumina, Cedrus Corp.). The NonSelective Response Mode consisted of either red and green Squares, or negative or neutral Faces stimuli randomized in a 50:50 distribution (100% response, 0% inhibition). The Selective Response Mode required responses selectively to either green Squares while inhibiting to red Squares, or to negative Faces while inhibiting to neutral Faces, randomized in a 80:20 distribution (80% response, 20% inhibition). Preceding the task runs, a non-task active baseline control condition was executed during which participants fixated on a black and white flashing checkerboard (8hz)^61^. All tasks (programmed using Presentation; Neurobehavioral Systems Inc.) were projected onto a screen positioned at the rear of the scanner for participants to view with an angled mirror attached to the head coil.

**Figure 1.**
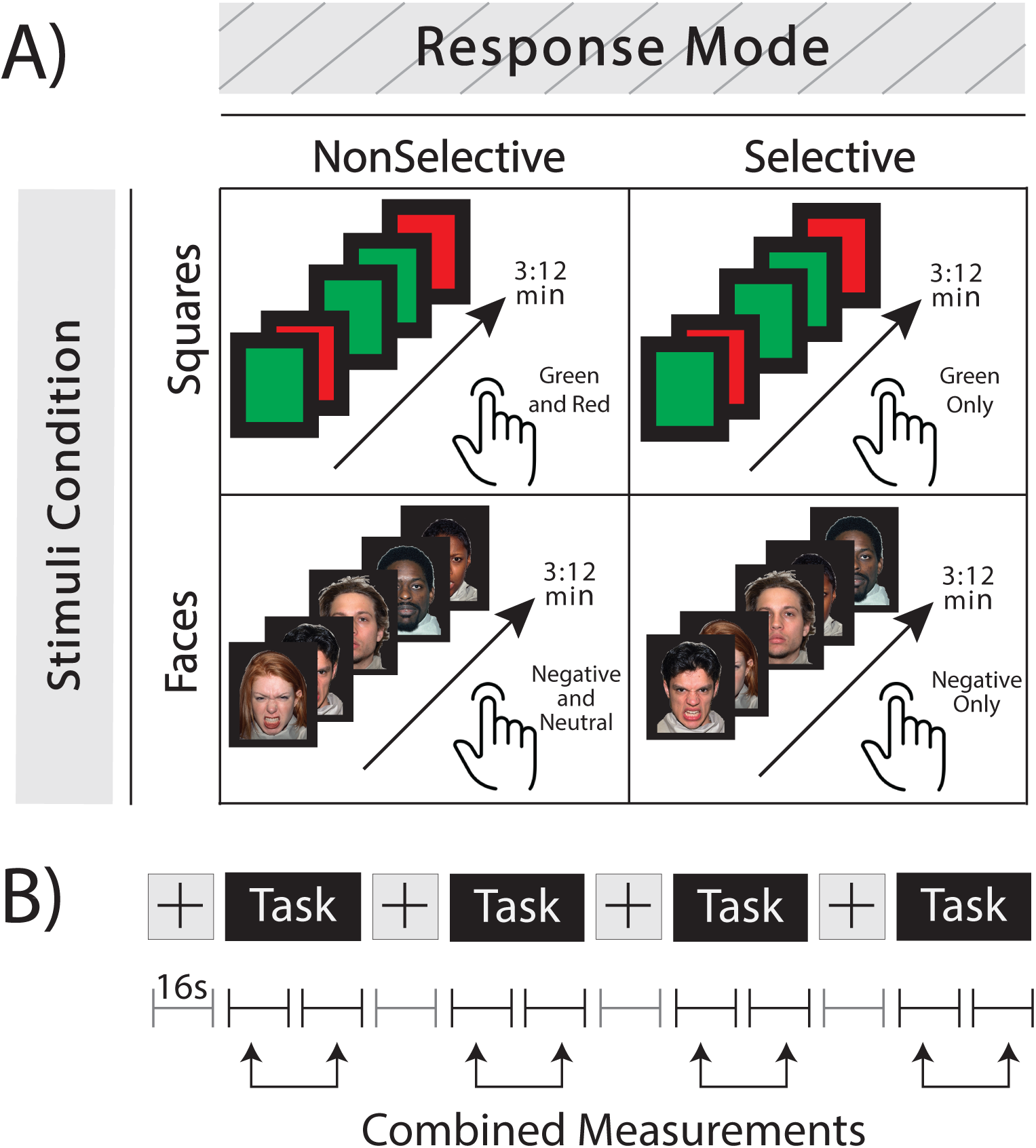
Schematic of the inhibitory motor control task design **A.** The Visually Guided Motor Response Inhibition Task consisting of 2 Response Modes, NonSelective and Selective, by 2 Stimuli Conditions, Squares and Faces. Participants respond to both cues in the NonSelective mode. Refer to the text for details on the instructions to participants. **B.** The timing of ^1^H MRS measurements per task run. Each run consists of four 32-second task blocks interleaved by 16-second rest blocks (fixation crosshair). Consecutive measurements within task blocks were averaged, yielding 4 separate outcome ¹H MRS measurements per task run.

Task performance for the Selective Response Modes, d-prime was computed, which reflects the standardized difference between the z-transformed hit rate and false alarm rate. Hit rate was defined as the proportion of correct taps to target stimuli, while false alarm rate was defined as the proportion of incorrect taps to non-target stimuli. For the NonSelective Response Modes, performance was assessed using hit rate. Additionally, the mean response time (RT) was evaluated across all blocks of each task run.

#### Neuroimaging Procedures

All scans were completed between 8 – 10am on a 3T Siemens MAGNETOM Verio^TM^ MRI system housed within the MR Research Facility, Wayne State University. The single-voxel point-resolved spectroscopy sequence (PRESS) was utilized to acquire the ¹H fMRS data from the midline dACC (20 x 17 x 12 mm^3^ or 4.08 ml). A T_1_-weighted set of MR images (multi-echo magnetization prepared rapid gradient echo sequence, ME-MPRAGE; TR= 2530ms; echo times = 1.79, 3.65, 5.51, 7.37ms; TI= 1100ms; flip angle = 7°; FOV= 256×256mm^2^; 176 slices; resolution= 1×1×1mm^3^, TA= 6:55) was first acquired to guide the placement of the ¹H MRS voxel, which encompassed the dorsocaudal BA 32 and BA 24. That is, the anatomical location and orientation of the ¹H MRS voxel was derived by co-registering a predefined voxel (in the MNI template brain) to the participant’s T_1_-weighted images using the auto-voxel placement procedure, which ensured consistent voxel locations between participants^62^. The PRESS sequence, which was developed by Edward J. Auerbach and Małgorzata Marjańska and provided by the University of Minnesota under a C2P agreement, included outer volume saturation (OVS) and water suppression by variable power and optimized relaxation delays (VAPOR)^63^. The acquisition parameters of the ¹H fMRS per task run included: TR= 4000ms, TE= 23ms, 4 averages per measurement, 16 second temporal resolution per measurement, 13 consecutive measurements, 2048 points, 2000Hz bandwidth and TA: 3:12min. The short-TE minimized signal attenuation and *J*-modulation effects, while the long repetition time minimized T_1_ relaxation partial saturation effects. FAST(EST)MAP shimming^64^ over the dACC voxel area was conducted prior to the data collection to minimize spectral linewidths, which were less than 8.75hz (**Table 1**). An unsuppressed water measurement was collected at the end of each scan run (TR: 10s, 4 averages and TA= 0:40min) and used for spectral processing and absolute quantification^65^. Lastly, to assess potential head movement between task run acquisitions, a set of localized images were acquired before and after each ¹H fMRS task run acquisition (TR= 8.6ms; TE= 4ms; FOV= 256×256mm^2^; 1 slice; resolution= 1×1×7mm^3^).

**Table 1.**
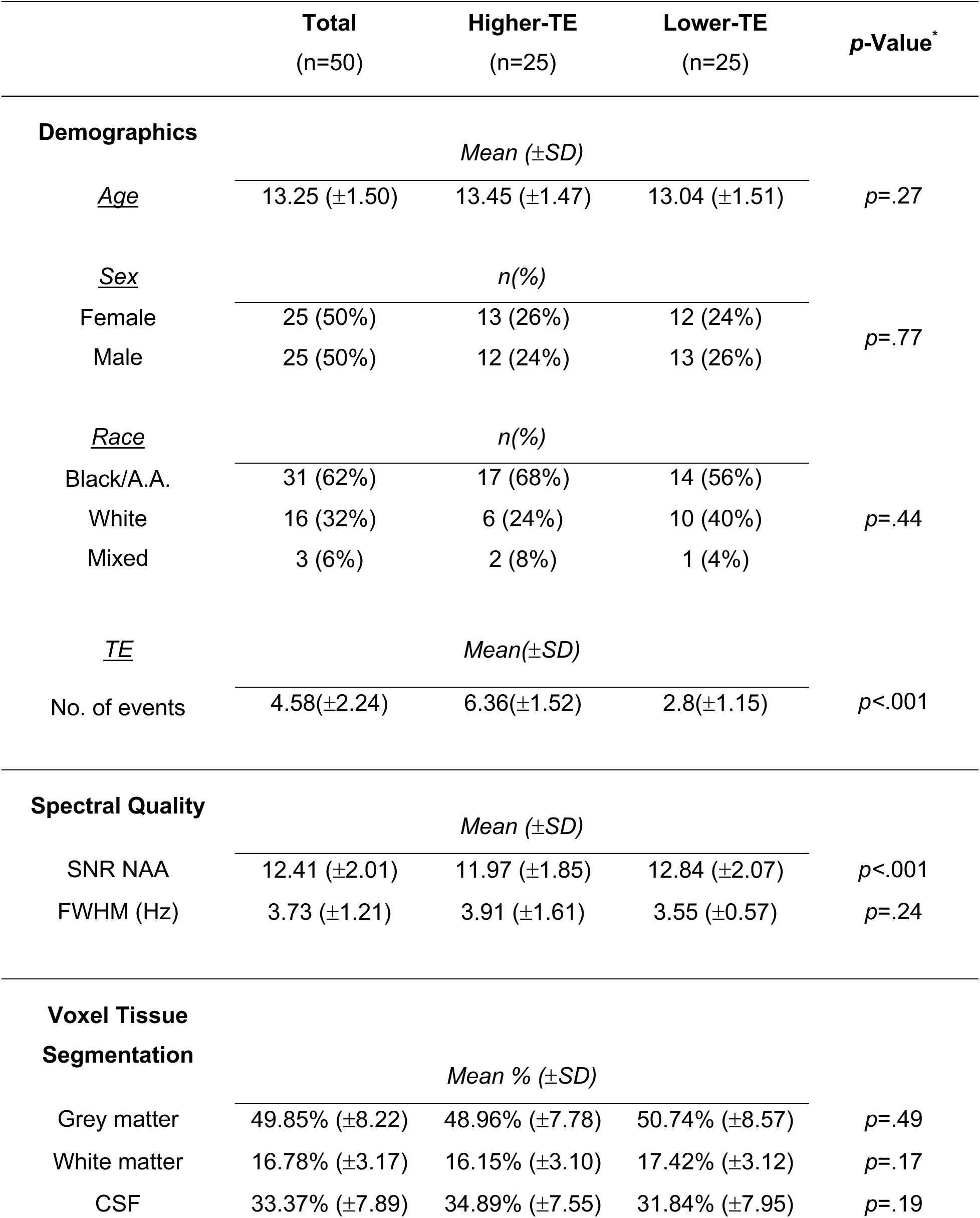
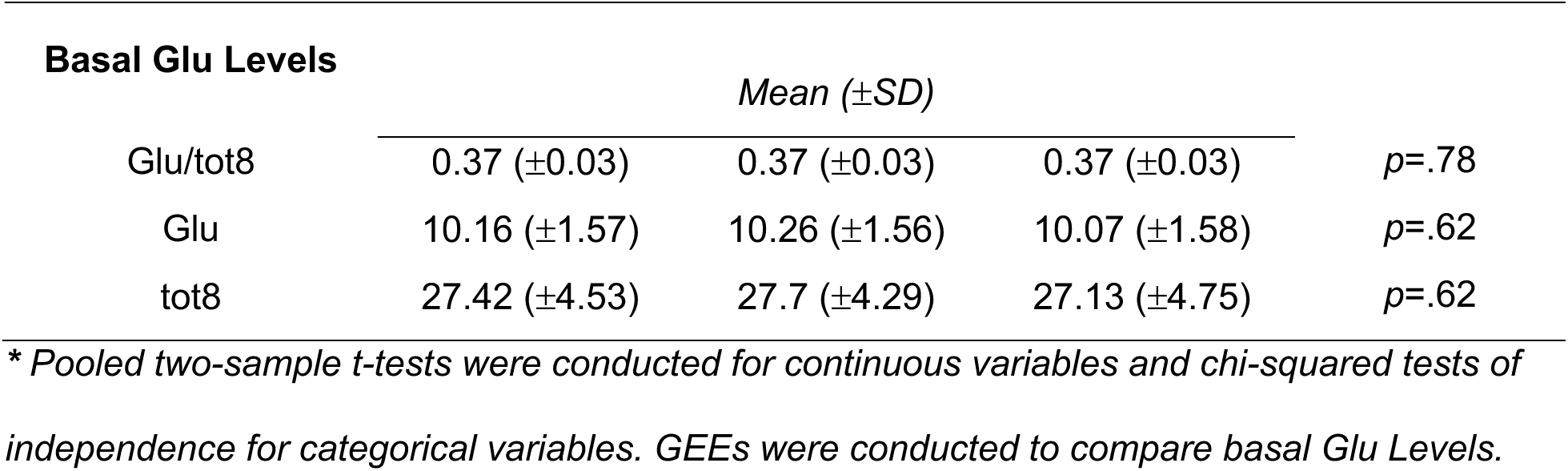
Comparison of High and Low TE-Groups.

Spectral processing and quantification procedure was fully automated and carried out using in-house scripts/programs and LCModel with a simulated basis set [N-acetyl aspartate (NAA), N-acetyl aspartyl glutamate (NAAG), phosphocreatine (PCr), creatine (Cr), glycerophosphocholine (GPC), phosphocholine (PC), myo-Inositol, glutamate, glutamine, GABA, lactate, glutathione (GSH), aspartate, alanine, glucose, taurine, scyllo-Inositol, and macromolecules/lipids]^66^. The first measurement of each task run was removed to allow steady-state equilibrium to be reached. The averaging of the consecutive ¹H MRS measurements within each task block was conducted using a two-step procedure. First, a mean 1^st^-order phase value was calculated by fitting the individual ¹H MRS measurements in each task run using the LCCORAW option. Second, each individual ¹H MRS measurements were re-fitted where the 0^th^-order phase was allowed to vary but the 1^st^-order phase was fixed to the derived mean value. The outcome re-fitted ¹H MRS spectra were phase and shift corrected and the two ¹H MRS spectra within each block were then averaged for final fitting (**Figure 2**), which yielded one Glu level estimate per block and hence, four per task run (**Figure 1b**). Moreover, a soft constraint was applied to the LCModel fitting by constraining the Glu to glutamine (Gln) ratio to reflect biologically plausible values (i.e., Glu/Gln = 4.0002 ± 2.6668)^67,68^. The imposed soft constraint on the Glu and Gln estimates did not affect the level of either metabolite but did significantly improve the reliability (i.e. reduce the coefficient of variation) of their estimated levels (**Supplement 1**). ¹H MRS spectra collected during the interspersed 16sec rest epochs were excluded from analyses. Tissue segmentation analyses were performed on the T_1_-weighted images using the ANAT tool from FSL and the estimated tissue fraction values within each voxel along with the unsuppressed water signal and relaxation values were utilized for absolute quantification^69,70^. To account for head movement-related variability in Glu estimates (described below) between task runs, Glu levels were expressed relative to the total ¹H amplitude signal of the eight highest non-Glu metabolites (termed tot8 and included NAA, NAAG, PCr+Cr, GPC+PC, myo-Inositol, glutamine, GABA and GSH), which served as the internal reference with the assumption that the relative Glu amplitude will scale approximately with the total signal amplitude if head movement occurred. The resulting Glu/tot8 ratio was used as the dependent variable in statistical analysis. The estimated spectral linewidth (FWHM) did not differ between the ¹H MRS spectra collected during the baseline control condition compared to task conditions (**Supplement 2**) and therefore, linewidth broadening correction for possible BOLD effects between conditions was not applied to the data^38^.

**Figure 2.**
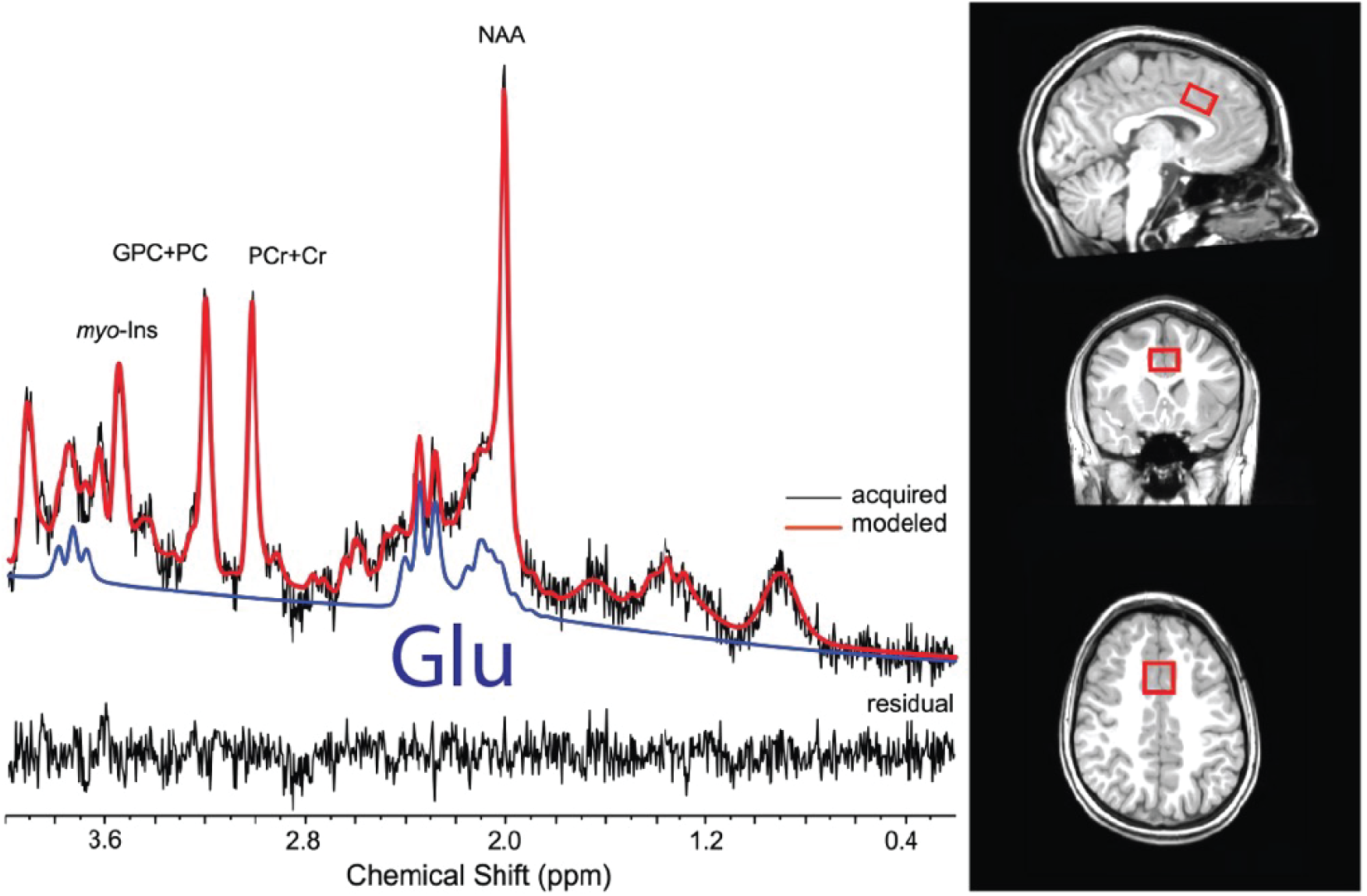
Example spectra and voxel placement A representative dACC voxel placement (right) and typical ^1^H MRS spectrum after combining two ¹H MRS measurements within a task block as outlined in Figure 1b.

Voxel placement accuracy and reliability was assessed by first co-registering the voxel location of each participant back to the MNI template space (via the inverse nonlinear transformation) followed by computing the percentage of overlapping pixels from each voxel with respect to the predefined template voxel^62^.

#### Spectral Data Quality Screening and Exclusion

Individual spectra were screened for: 1) insufficient quality, 2) potential head movement, and/or 3) poor task compliance. Insufficient quality was defined as any ¹H MRS spectra with a S/N (of the NAA signal) less than 5, a FWHM > 10 Hz, a Cramér-Rao Lower Bound (CRLB) value from the main metabolites (Glu, NAA, GPC+PC, PCr+Cr, and myo-Inositol) >20%, or with a CRLB for Gln >35%. Any head movement between consecutive ¹H MRS measurements will lead to shifts in the anatomical location of the localized ¹H MRS voxel, which in turn may have an impact on the shimming condition (FWHM) and signal amplitude of metabolites (S/N of NAA and total signal amplitude). Therefore, potential head movement was assessed excluding ¹H MRS spectra demonstrating substantial changes (>10%) in the FWHM, S/N and/or the total signal amplitude of the 8 main metabolites relative to the ¹H MRS spectra acquired during the baseline control condition. Additionally, we corroborated potential head movement qualitatively by visualizing overlaid MRI images from the localizer scans acquired between each task run and preceding images to screen for head movement between task runs (**Supplement 3**). Poor task compliance was defined as ≤50% task accuracy for a set of trials within a task block. If half or more of a participant’s data was removed for any reason listed above (i.e. 2 or more task runs), the participant was excluded from statistical analysis. All data from 3 participants were excluded, resulting a final sample of 50 participants.

## Statistical Analyses

Repeated measures generalized estimating equations (GEE) analyses tested significant changes in Glu/tot8 and task performance metrics across task conditions, using a normal distribution with an identity link, an exchangeable working correlation structure, and clustering within participants (SAS GENMOD, SAS Institute Inc). For hypothesis one, a full factorial GEE was used to examine the three-way interaction among TE-Group, Stimuli Condition, and the Response Mode terms. Additionally, the mean Glu/tot8 ratio acquired during the checkerboard baseline control condition was added as a covariate to reflect the zero-modulation reference point. Given the relatively limited sample size, a two-tiered analytic approach was also employed: (1) Response Modes were initially collapsed to test the TE-Group by Stimuli Condition interaction, and (2) follow-up models examined the TE-Group by Response Mode interaction within each level of Stimuli Condition. Post hoc comparisons were conducted using Least Squares Means comparisons. For hypothesis 2, GEEs tested associations between Glu/tot8 modulation during Faces and anxiety/PTSD symptoms, controlling for age and sex. Supplemental analyses included an additional full factorial GEE examining the effect of TE as a continuous variable, indexing the degree of TE, to examine a potential dose-dependent relationship with the outcome variables.

Statistical significance was determined at α = 0.05. An a priori power analysis was conducted using G*Power^71^ to estimate statistical power for a sample size of *n* = 50. Assuming a small-to-moderate effect size (Cohen’s *d* = 0.25) and an alpha level of 0.05, the analysis indicated 80% power to detect a between-group difference.

## Results

### Demographics, Spectral Quality, and Voxel Placement

A median-split based on TE was applied to categorize the sample into two groups: the Higher TE-Group (6±1events) and Lower TE-Group: (3±1events), **Table 1**. The groups did not differ significantly in age, sex, race, tissue fraction of grey and white matter within the ¹H MRS voxels, basal levels of Glu/tot8 collected during the non-task checkerboard control condition (*ps*>0.05; **Table 1**), or voxel placement accuracy (percent overlap relative to the template voxel) [*t*(51)=0.32, *p*=.789; **Figure 3**]. The Higher TE-Group exhibited lower S/N of the NAA signal than the Lower TE-Group [*t*(706)=-5.86, *p*<.001; **Table 1**]. Following spectral quality assessment, 9.4% of observations (94/1000) were excluded, and the mean CRLB for Glu across the final dataset was 5.9 ± 0.8 (range=4-9).

**Figure 3.**
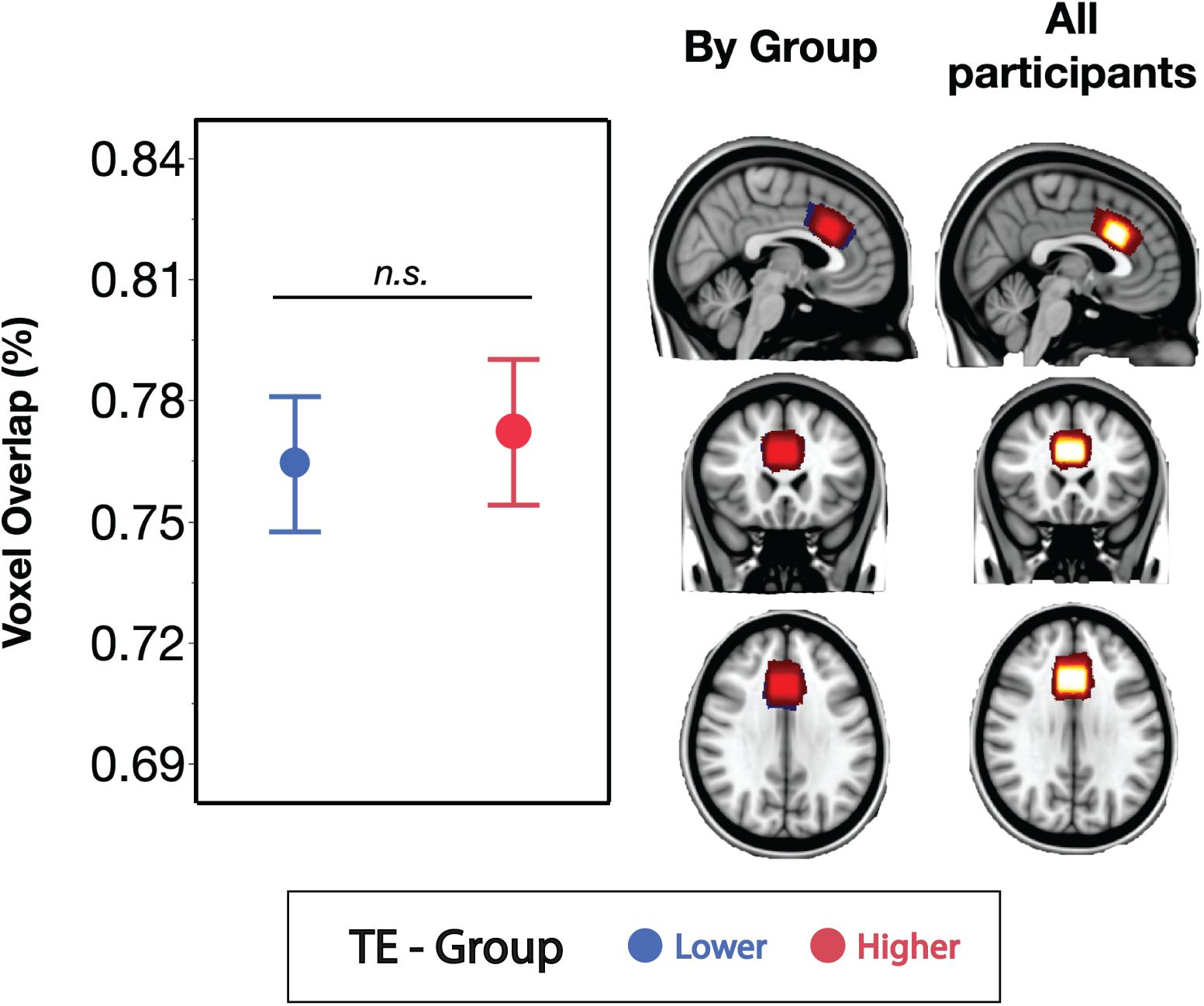
Voxel overlap with an *a priori* defined dACC template voxel **Left:** Voxel placement accuracy, quantified as the percentage of overlap with the template voxel, did not differ significantly between TE-Groups (*t*(51)=0.32, *p*=.789). **Right:** Visualization of voxel placement consistency between TE-Groups (red=Higher; blue=Lower) and across all participants (red-white color gradient, with white indicating 100% voxel overlap across participants).

### dACC Glu/Tot8 Modulation

In the full-factorial model with the three terms, TE-Group, Stimuli Condition (Squares vs Faces), and Response Mode (Selective vs NonSelective), the three-way interaction term and the covariate was non-significant (χ² = 0.17, *p* =.684). Consequently, adopting a two-step approach firstly revealed a significant Stimuli Condition by TE-Group interaction (χ^2^=4.66, *p*=.031; **Figure 4a**). Post-hoc analysis revealed that the Higher TE-Group demonstrated a significantly lower Glu/tot8 compared to the Lower TE-Group during Faces (z=-2.28, *p*=.023), but not during Squares (*z*=-.22, *p*=.82). Also, the Higher TE-Group demonstrated a significant within-group reduction in

**Figure 4.**
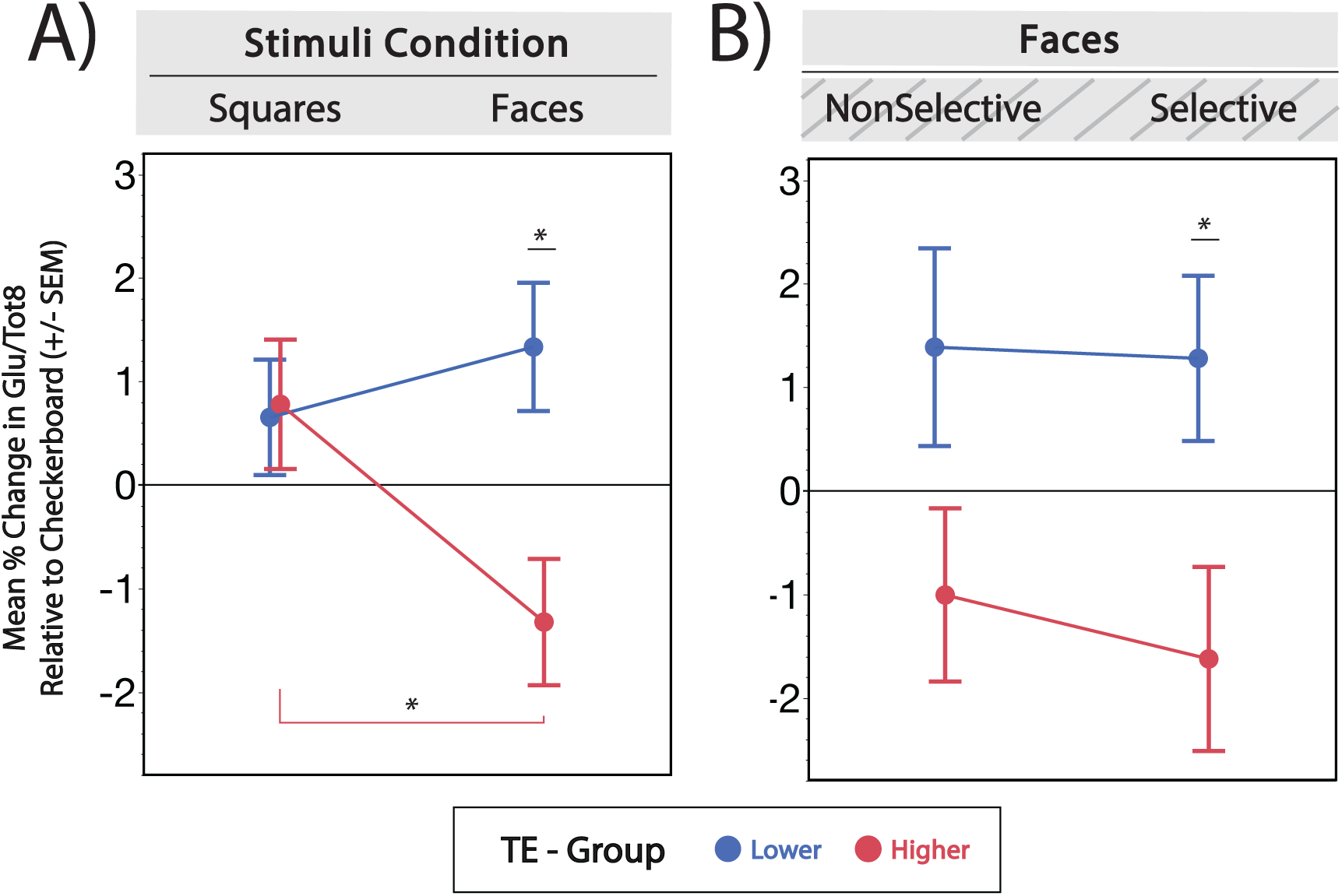
Glu Modulation The mean ± standard error (error bars) percent change in Glu/tot8 relative to the task-inactive checkerboard baseline by task run and TE-Group. **A.** Significant TE-Group by Stimuli Condition interaction (χ^2^=4.66, *p*=.031). Post-hoc mean comparisons demonstrated a significant difference in Glu/tot8 between TE-Groups during Faces (z=-2.28, *p*=.023), but not Squares (*z*=-.22, *p*=.82). The Higher-TE group also demonstrated a significant reduction in Glu/tot8 from Squares to Faces (*z*=3.39, p<.001) while the Lower-TE group showed no change (*z*=-0.71, *p*=.476). **B.** Within the Faces Stimuli Condition, there was a significant main effect of TE-Group (χ^2^=4.33, p=.037). Post hoc analyses revealed a significant TE-Group difference during Selective responding (z=-2.06, p=.040) but not NonSelective (z=-1.44, p=.150). Exploratory post-hoc analysis showed a significant TE-Group difference during the Selective (*z*=-2.13, *p*=.03) but not the NonSelective Response Mode (*z*=-1.44, *p*=.15).

Glu/tot8 during Squares compared to during Faces (*z*=3.39, p<.001), whereas the Lower TE-Group did not significantly differ between Conditions (*z*=-0.71, *p*=.476). We additionally tested the Stimuli Condition by TE (as a continuous variable) interaction, which was also significant (χ^2^=7.94, *p*=.005; **Supplement 4**).

In our second step, we tested if the TE-Group difference observed during the Faces Stimuli Condition was specific to Selective or NonSelective responding. Within the Faces Stimuli Condition, the Response Mode by TE-Group interaction was not significant (χ^2^=0.18, p=.670). However, the TE-Group term was significant (χ^2^=4.33, p=.037) such that, on average, the Higher-TE group demonstrated lower Glu/tot8 compared to the Lower TE-Group across both response modes (**Figure 4b**). Within the Squares Stimuli Condition, the main effect of TE-Group and the Response Mode by TE-Group interaction were not significant (ps>0.05; **Supplement 5**).

### Behavioral Performance

#### Hit Rate

During NonSelective responding, there were no significant main effects of TE-Group (χ^2^=0.01, *p*=.90) and Stimuli Condition (χ^2^=0.91, *p*=.34), or TE-Group x Stimuli Condition interaction effects (χ^2^=0.99, *p*=.32; **Figure 5a**).

**Figure 5.**
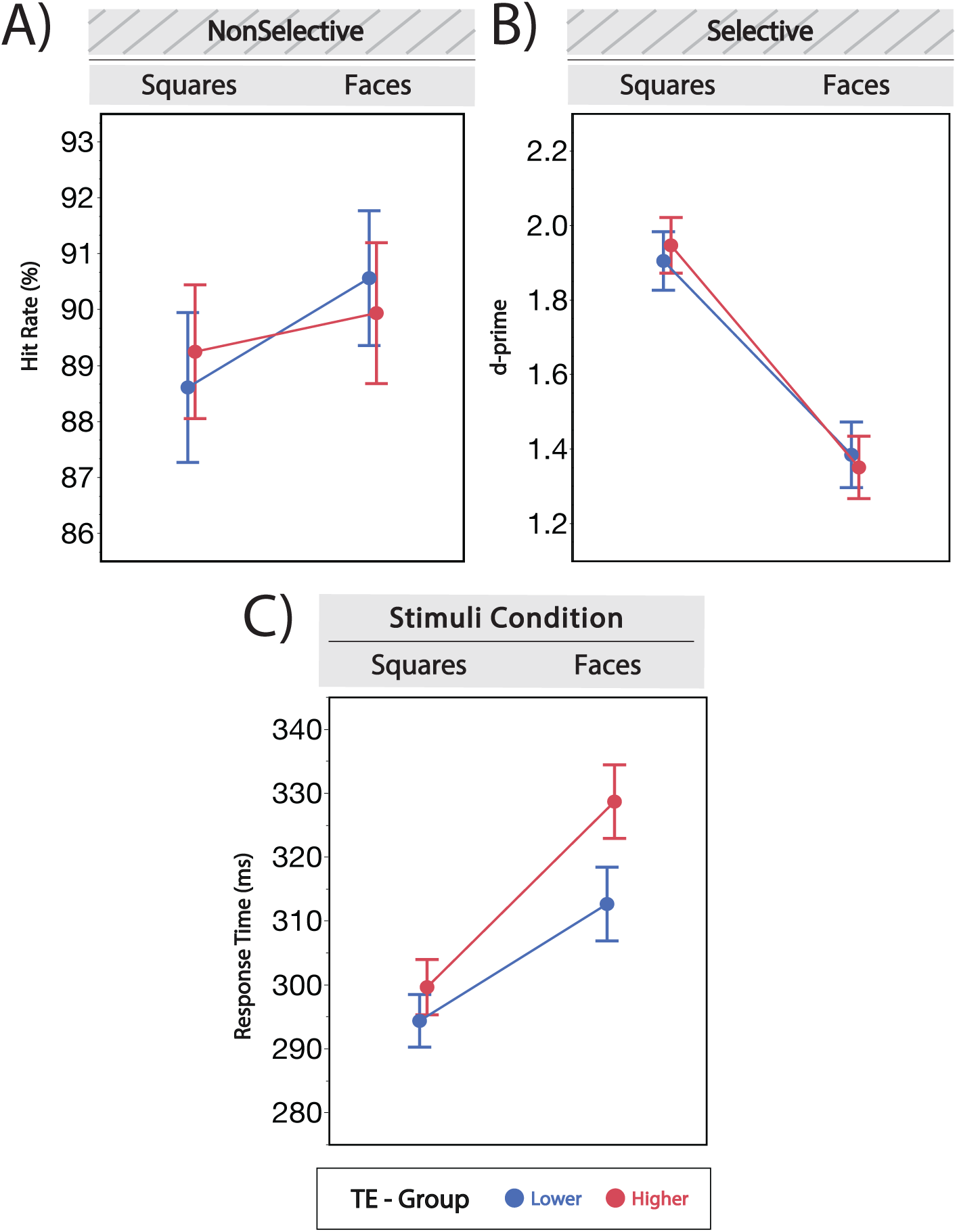
Behavioral performance The mean ± standard error (error bars) behavioral performance metrics by task run and TE-Group. **A.** During the NonSelective Response Mode, hit rate did not show a significant effect of Stimuli Condition, TE-Group, nor a Stimuli Condition x TE-Group interaction (*ps*>0.05). **B.** During the Selective Response Mode, d-prime showed a significant main effect of Stimuli Condition (χ^2^=26.88, *p*=.005) **C.** Across the full task, response time showed a trend-level interaction between Stimuli Condition and TE-Group (χ^2^=2.91, *p*=.088).

#### D-Prime

During Selective responding, the interaction between TE-Group and Stimuli Condition was non-significant (χ^2^=0.72, p=.395) as well as the main effect of TE-Group (χ^2^=0.00, *p*=.969). The main effect of Stimuli Condition was significant, (χ^2^=26.88, *p*=.005) such that participants demonstrated smaller d-prime values for Faces relative to Squares independent of TE-Group (**Figure 5b**).

#### Response Time

The full factorial model showed no significant three-way interaction among TE-Group, Stimuli Condition, and Response Mode (χ^2^=1.50, *p*=.221; **Supplement 6**). The Stimuli Condition x TE-Group interaction was also not significant (χ^2^=2.91, *p*=.088; **Figure 5c**), however, the Stimuli Condition x Response Mode interaction was significant (χ^2^=14.70, *p*<.001), such that participants demonstrated significantly longer RTs for Faces relative to Squares during the Selective Response Mode (χ^2^=5.98, *p*<.001), but not the NonSelective Response Mode (χ^2^=4.50, *p*=.144).

#### Associations with Anxiety Symptom Severity

The multiple linear regression model examining the relationship between Glu/tot8 modulation observed during the Faces Stimuli Condition and anxiety symptom severity controlling for age and sex was non-significant (*F*(3,46)=1.29, *p*=0.851; *R^2^*=.02). None of the individual predictors reached statistical significance (*ps*>0.05). Similarly, hit rate, D-prime, and response time did not significantly predict anxiety symptoms when controlling for age and sex (*ps*>0.05).

## Discussion

The present study examined differences in dACC Glu modulation between adolescents with lower and higher TE (*Mdn*=4 events). Glu modulation was assessed relative to a checkerboard baseline condition (expressed as a ratio to the tot8 signal) during an inhibitory motor control task with and without emotional stimuli interference, using ¹H fMRS. TE-Groups differed in Glu modulation during task performance with emotional Face stimuli, but not with non-emotional Square stimuli. Specifically, youth with Higher TE demonstrated significantly lower dACC Glu modulation relative to the Lower TE-Group, during the Faces Stimuli Condition (**Figure 4a**). Additionally, the Higher TE-Group showed a significant reduction in Glu/tot8 from the Squares to Faces condition resulting in negative Glu modulation, whereas the lower TE-Group maintained positive modulation (**Figure 4a**). Consistent with this, supplemental analysis treating TE as a continuous variable showed that greater TE was associated with larger magnitude decreases in dACC Glu during the Faces condition (**Supplement 4**). There was no significant group difference during the Squares Condition (**Supplement 5**). We conclude that youth with a greater degree of TE demonstrate less Glu modulation during task performance with emotional stimuli relative to youth with less exposure, suggesting heightened TE-related vulnerability to emotion interference with dACC Glu responses during inhibitory control.

Next, we tested if this emotion interference effect (i.e. the TE-related difference in Glu modulation during Faces) was specific to the task condition involving inhibitory control, or the Selective Response Mode. Counter to our hypothesis, we found no TE-Group x Response Mode interaction during the Faces Stimuli Condition, suggesting that the group difference in Glu modulation was not specific to inhibitory control. The Higher TE-Group demonstrated lower modulation on average across both Response Modes. This finding suggests that the dACC may have been engaged by stimulus salience and/or complexity, consistent with prior fMRI evidence of its role in salience detection in trauma-exposed populations^72,73^. Future studies should examine dACC specificity for cognitive processes and/or emotional reactivity in TE youth. Taken together, these findings suggest that youth with greater TE are more susceptible to emotion interference of cognitive control more generally. Importantly, the mean dACC Glu level and Glu/tot8 ratio acquired during the checkerboard control condition, which may reflect the “basal” metabolic state, did not differ between TE-Groups (**Table 1**). This stresses the importance of utilizing ¹H fMRS over conventional ¹H MRS to detect task-related neurometabolic changes associated with psychiatric risk factors such as TE.

We also found no significant group differences in task accuracy, measured with hit rate and d-prime, suggesting the groups did not differ in task engagement. While not significant, the trending Stimuli Condition x TE-Group interaction when testing response times (**Figure 5c**) is consistent with prior literature demonstrating that TE youth are disproportionately impacted by emotion interference of inhibitory control when assessing task performance using an emotional go/no-go paradigm^8^. Additionally, we found a significant interaction between Stimuli Condition and Response Mode for response times, such that all participants on average demonstrated significantly slower response times for Faces relative to Squares under the Selective Response Mode when inhibitory control was required. This supported the validity of the task in eliciting the expected cognitive control processes.

Reasoning that the tot8 signal amplitude does not modulate under task conditions, then the significant group differences in the Glu/tot8 modulation can be interpreted to reflect temporal changes in Glu levels within the dACC under varying task conditions. Task-evoked changes in Glu levels can be interpreted within the framework of dynamic excitatory and inhibitory (E/I) balance, which posits that neurotransmission drives energy metabolism within the cortex to achieve varying steady states of excitatory and inhibitory neurotransmission subserving cognitive function^46^. Within this framework, Glu modulation detected by ^1^H fMRS reflects net changes in excitatory neurotransmission averaged across microcircuits within the voxel and over the acquisition period. Further, increased engagement of excitatory neuronal populations is accompanied by a net increase in neurometabolic flux leading to Glu synthesis^36,39,40^. As such, the observed TE-Group differences in Glu modulation during the Faces Stimuli Conditions may reflect variability in the capacity to dynamically shift E/I balance across microcircuits within the dACC during emotion interference of cognitive control. Specifically, the net negative shift in E/I balance observed within the Higher TE-Group from Squares to Faces may reflect a net shift in E/I balance away from excitatory neural activity, accompanied by diminished synthesis of Glu across dACC microcircuits observed within the ^1^H fMRS voxel during these task conditions.

One possible explanation for the current study’s observation of TE-related Glu/tot8 modulation differences is that youth with higher TE experience greater interference from emotionally salient stimuli, biasing neural resources toward bottom-up emotional processing and away from top-down cognitive control. During the Faces Stimuli Condition, negative emotional stimuli may have contributed to altered dACC glutamatergic responses. Although the present findings cannot directly address mechanism, TE may alter excitatory and inhibitory microcircuits within stress susceptible regions like the dACC. Evidence from animal models demonstrates that stress exposure (e.g., restraint stress) induces structural remodeling of the dendritic arbors and spines of excitatory pyramidal cells in the medial PFC^74,75^, likely partially mediated through corticosterone release via the hypothalamic-pituitary-adrenal (HPA) axis^76^. These stress-induced changes are associated with lasting alternations in microcircuit connectivity^77^. In addition to excitatory neurons, trauma exposure also affects cortical inhibitory neurons. During adolescence in animal models, stress has been shown to reduce the number of mPFC parvalbumin-positive (PV+) interneurons and density of perineuronal nets (PNNs) that surround these cells and regulate their activity^78,79^. Together, these findings suggest that TE-related alterations in cortical microcircuit organization could contribute to differences in dynamic E/I balance shifts exploited under specific emotional contexts. However, this interpretation requires further investigation in future studies.

We further hypothesized that Glu modulation observed during the Faces Stimuli Condition would predict anxiety symptom severity. Contrary to our hypothesis, we found no significant association between Glu modulation during Faces and anxiety. The null finding may reflect a restricted range of anxiety symptoms in the current study, who endorsed fewer anxiety symptoms compared to another investigation of adolescents using the same measure^80^. Some participants may not yet have developed clinically meaningful anxiety symptoms given the typical age of onset for anxiety disorders^81^. The present study also excluded adolescents diagnosed with an anxiety disorder to ensure TE-related effects were not driven by pre-existing clinical symptoms, further limiting the range of symptoms. These factors may have limited variability necessary to detect a relationship between Glu modulation and anxiety. Future research should implement prospective, longitudinal designs to determine if TE-related differences in cortical Glu modulation during emotion interference of cognitive control in youth predicts the emergence of anxiety symptoms across a broader adolescent age range.

The present study has several limitations. First, the Stimuli Condition model term contrasted Squares vs Face stimuli, potentially confounding the effect of emotion with visual complexity and social vs non-social cues. Still, this is among the first studies to identify stimuli driven differences in Glu modulation in youth with trauma exposure, providing a foundation for future work. As emotion interference is typically linked to negative stimuli^8,82,83^, future ^1^H fMRS investigations should test specificity in Glu modulation to negative emotional stimuli in TE-youth. Second, ^1^H fMRS captures signal from all mobile Glu molecules within the voxel, and not synaptic Glu specifically. However, given the close link between oxidative metabolism and neurotransmission^46,84^, Glu modulation likely reflects, in part, task-evoked neurometabolic activity.

To our knowledge, this is the first study to examine TE-related differences in cortical Glu modulation using ^1^H fMRS during emotion interference of cognitive control in adolescents. These findings suggest that higher trauma exposure is associated with less prefrontal Glu modulation during emotionally salient task conditions, indicating altered excitatory neurotransmission during cognitive and emotional integration. Future research may further elucidate the neurometabolic pathways underlying excitatory neurotransmission in TE youth, potentially identifying novel biomarkers and therapeutic targets for trauma-related mental illness.

## Acknowledgements

This work was supported by the National Institute of Mental Health (Grant Nos. R01MH111682 and R01MH100122 [PI: TJ] and F31MH132307 [PI: JMF]). The authors thank the participants and their families for their time and commitment to this research. We also acknowledge the staff of the Wayne State University MR Research Facility (MRRF) for their assistance.

## Supplemental Material

**Supplement 1.** Glu/Gln Soft Constraints To assess the impact of imposing a soft constraint (S.C.) between Glu and Gln prior to LC model fitting, we used pairwise comparisons to assess differences in the mean steady state level and the coefficient of variation (CV) of Glu and Gln with and without the soft constraint imposed. We found that imposing the Glu/Gln soft constraint maintained the absolution Glu/Gln concentration ratio to be within physiological range (**Figure S1a**). Additionally, imposing the constraint did significant change Glu nor Gln concentrations (Glu: *t*(1419) = 0.76, *p* =.45; Gln: *t*(1419) =-0.17, *p* =.87) (**Figure S1b**) but did significantly improve the CV, or stability of their measurement (Glu: *t*(1419) =-2.7, *p* <.01; Gln: *t*(1419) =-5.9, *p* <.001) (**Figure S1c**).

**Figure S1.**
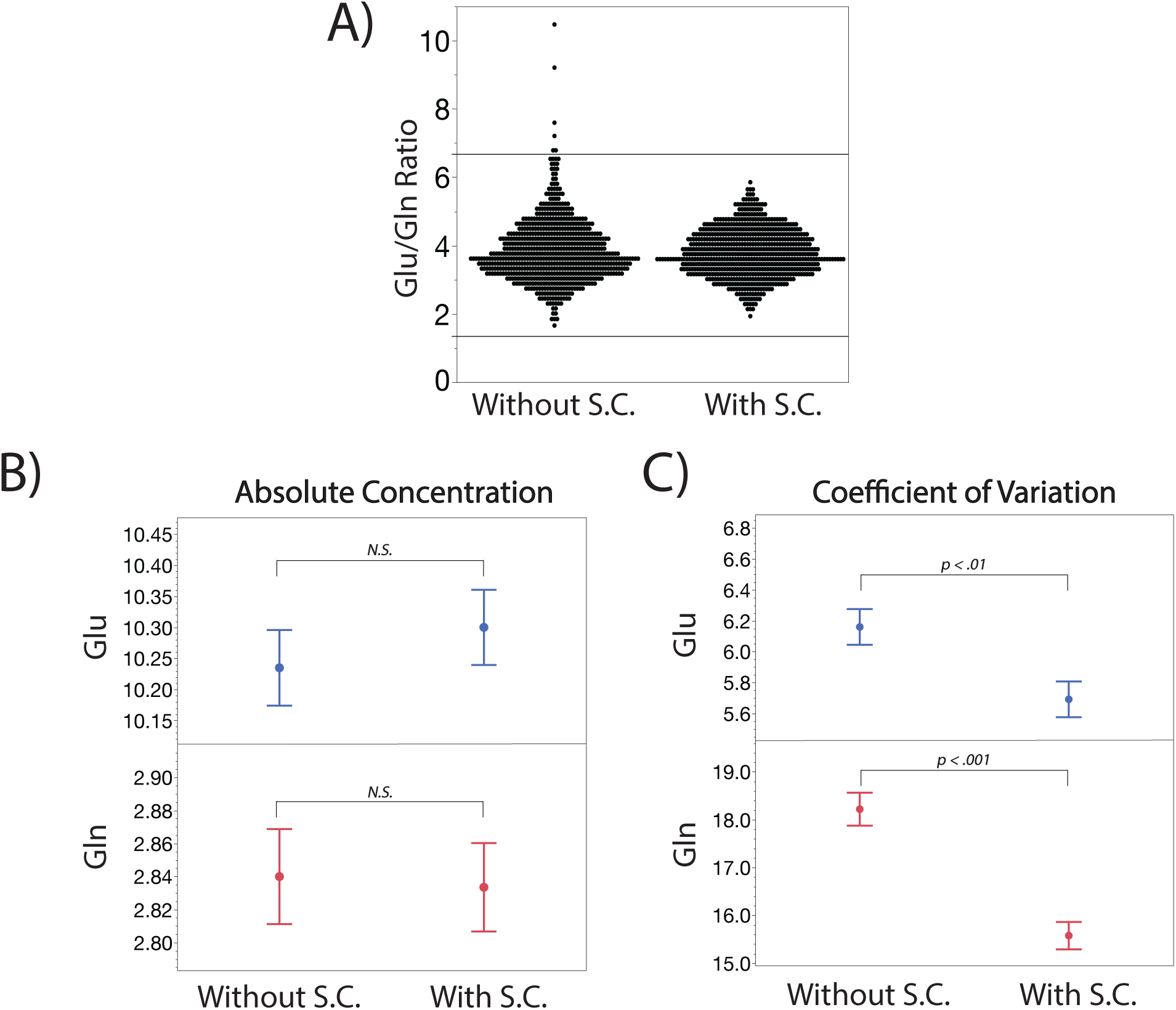
Glu/Gln Soft Constraints **A.** Imposing the Glu/Gln soft constraint (S.C.) maintained Glu/Gln within physiological range (i.e., 4.0002 ± 2.6667, top and bottom y-axis lines). **B.** Imposing the Glu/Gln S.C. did not significantly alter Glu and Gln concentrations collected during task runs (*ps >* 0.05). **C.** Glu/Gln S.C. significantly improved the coefficient of variation of Glu (*t* =-2.7, *p <* 0.01) and Gln estimates collected during task runs (*t* =-5.9, *p <* 0.001).

**Supplement 2.** BOLD Effects To examine potential BOLD effects, we utilized a repeated measures GEE approach to assess mean differences in FWHM between the baseline flashing checkerboard condition and the task runs. We repeated this analyses to test the effects of Stimuli Conditions and the Response Modes, similarly to our statistical approach for hypothesis testing. We found no significant main effect of Stimuli Condition (χ^2^=1.20, *p*=0.55), **Figure S2a**, nor Response Mode (χ^2^=2.15, *p*=0.34), **Figure S2b**, indicating BOLD effects were absent from our hypothesis tests.

**Figure S2.**
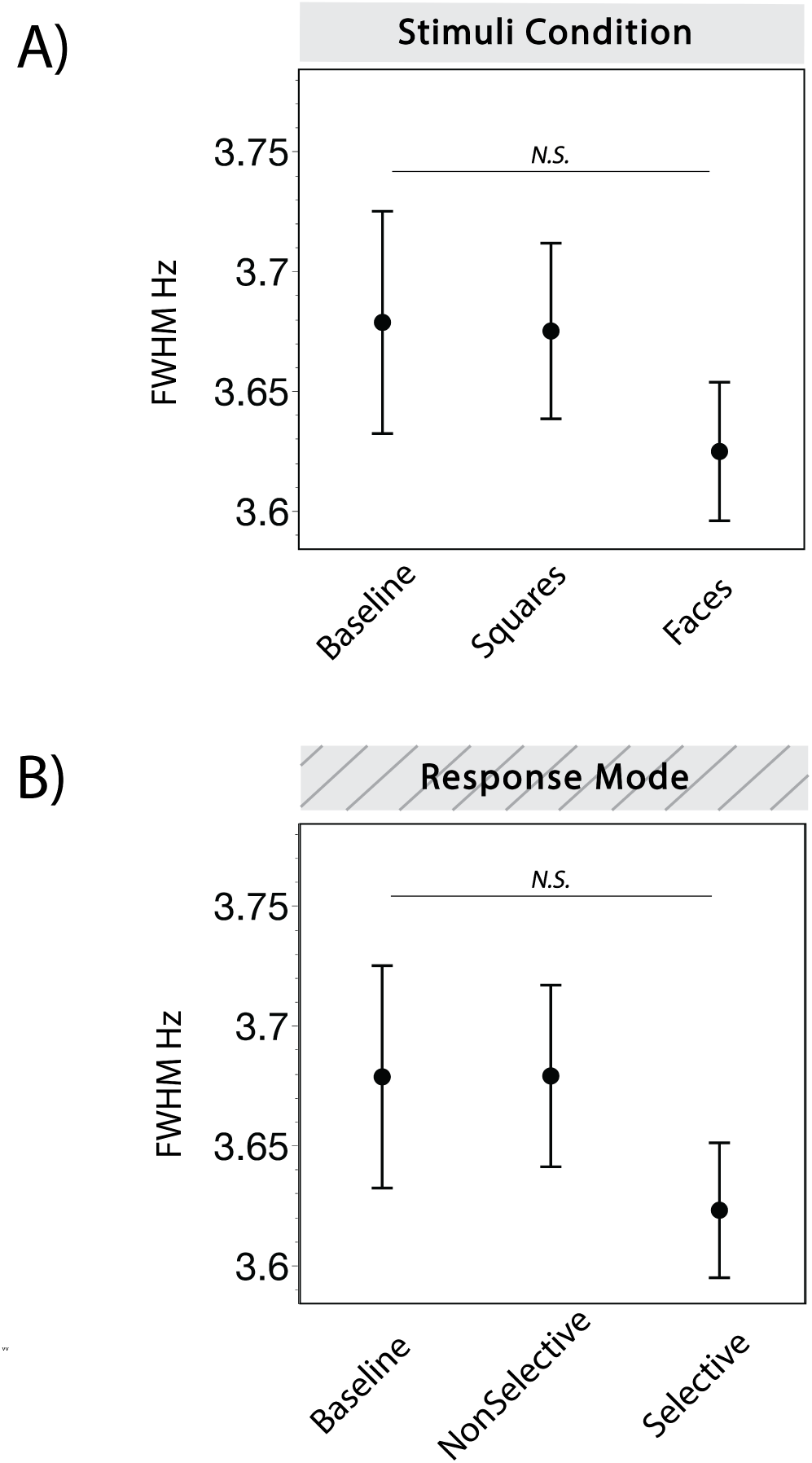
BOLD Effects **A.** No significant differences in FWHM between baseline and task runs split by Stimuli Condition (χ^2^= 1.20, *p* = 0.55). **B.** No significant differences in FWHM between baseline and task runs split by Response Mode (χ^2^= 2.15, *p* = 0.34).

**Supplement 3.** Participant Head Movement Assessment The task protocol, which includes multiple task run acquisitions, is prone to potential head movement effects especially in this young sample. Therefore, participant head movement between task runs was assessed using an in-house script designed to overlay each set of localized images with respect to the 1^st^ set acquired prior to the baseline control acquisition, which served as the reference for the anatomical placement of the dACC voxel location. The script generated a set of GIF images to visualize: (1) *intra-run head movements* by overlaying localizers collected before and after each ¹H MRS task run and (2) the *accuracy of voxel placement* throughout the course of the scan relative to the reference set of localizers. Using this output, head movement suspects were evaluated by visualization.

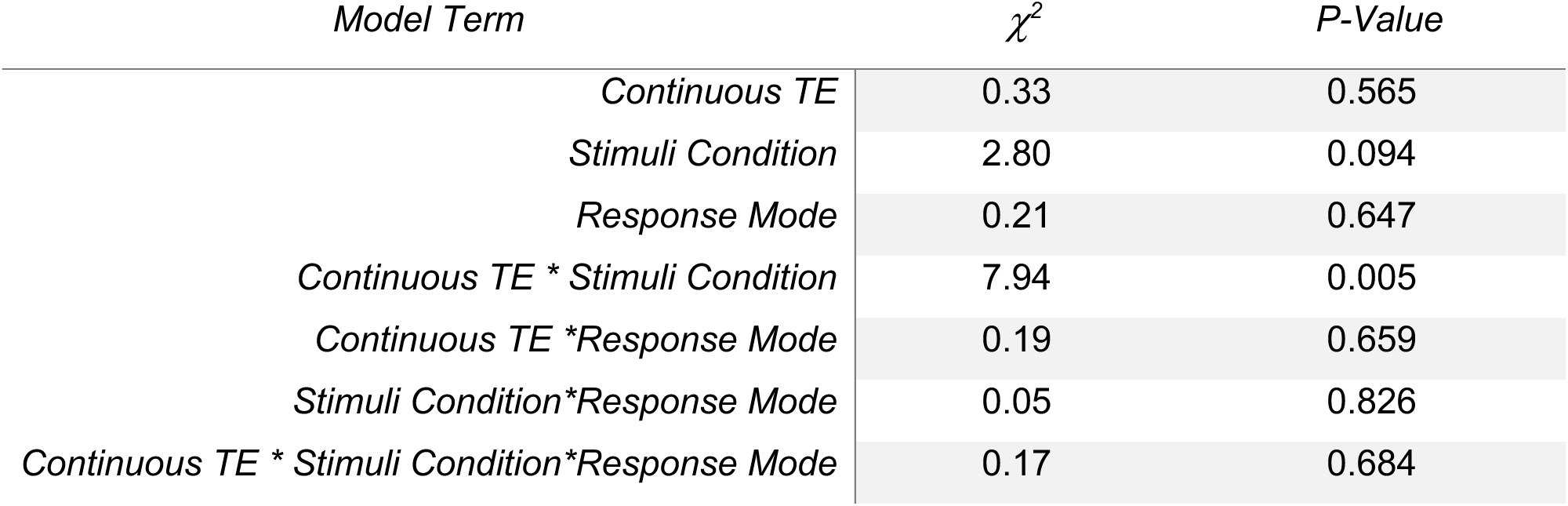

**Figure S4.**
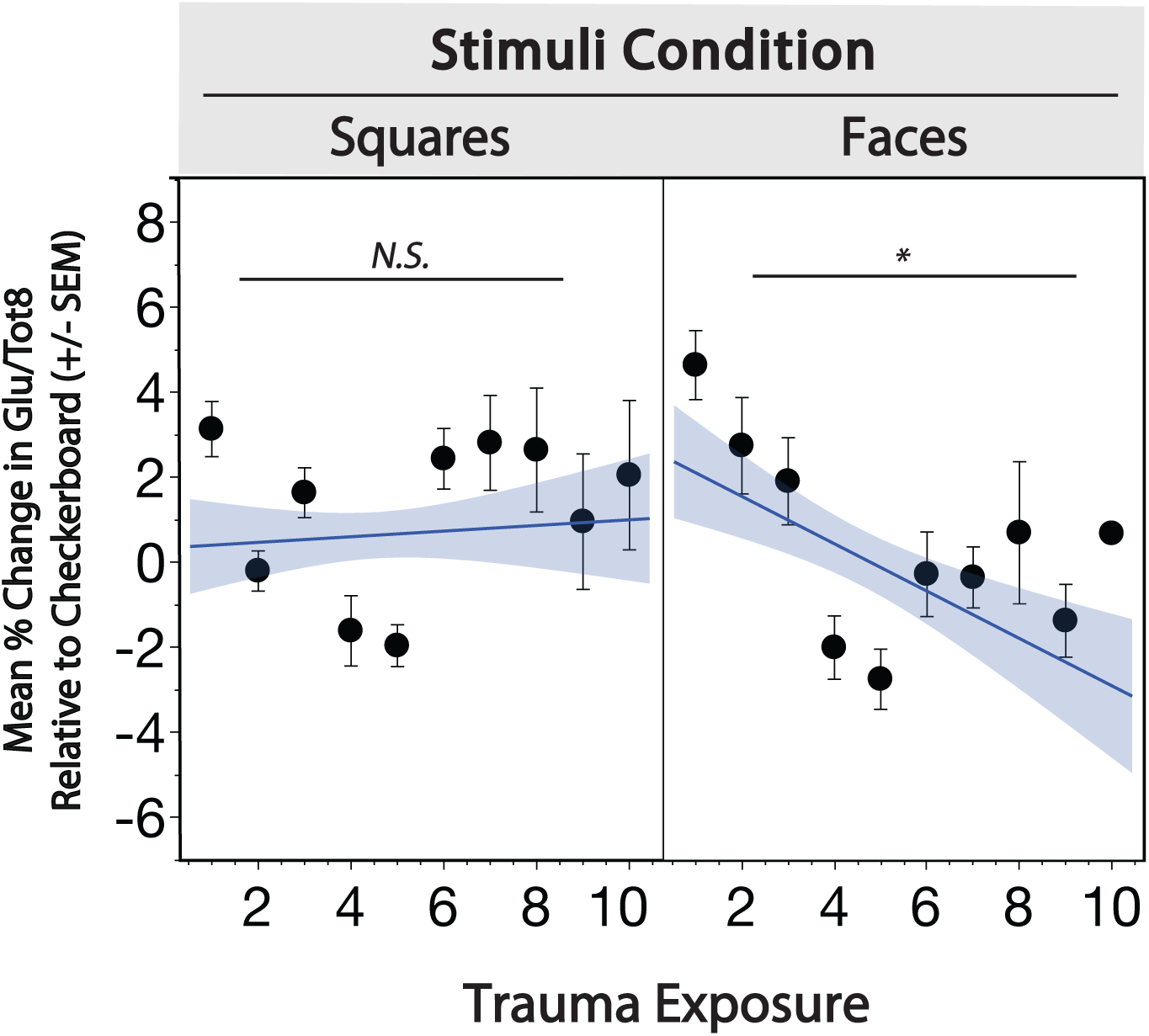
Continuous TE Pierson Correlations between Glu/tot8 modulation and continuous TE split by Stimuli Condition and collapsed across Response Mode. No significant association during Squares (*r^2^*=0.001, *p*=.591). For Faces, there is significant negative association between Glu/tot8 modulation and continuous TE (*r^2^*=0.041, *p*<.001).

**Supplement 4.** Continuous TE The full factorial GEE model including continuous TE as a continuous predictor of Glu/tot8 Modulation indicated a significant interaction between continuous TE and Stimuli Condition (χ^2^=7.94, *p*=0.005). The main effects of continuous TE and Stimuli Condition were non-significant (*ps*>0.05). **Figure S3** demonstrates the associations between continuous TE and Glu/tot8 by Stimuli Condition.Pierson Correlations between Glu/tot8 modulation and continuous TE split by Stimuli Condition and collapsed across Response Mode. No significant association during Squares (*r2*=0.001, *p*=.591). For Faces, there is significant negative association between Glu/tot8 modulation and continuous TE (*r2*=0.041, *p*<.001).

**Supplement 5.** Squares Stimuli Condition, Glu/Tot8 Within the Squares Stimuli Condition, a GEE model was used to examine the effect of Response Mode and its interaction with TE-Group on Glu/tot8 modulation during the Inhibitory Motor Control task.

| *Model Term* | $\chi^2$ | *P-Value* |
| --- | --- | --- |
| *TE-Group* | 0.01 | 0.913 |
| *Response Mode* | 1.19 | 0.276 |
| *TE-Group\*Response Mode* | 0.09 | 0.760 |

**Supplement 6.** Full Factorial GEE, Response Time To assess the effect of a three-way interaction among TE-Group, Stimuli Condition, and Response Mode, a GEE model was used to predict Response Time during the Inhibitory Motor Control task.

| *Model Term* | $\chi^2$ | *P-Value* |
| --- | --- | --- |
| *TE-Group* | 0.61 | 0.436 |
| *Stimuli Condition* | 20.44 | <0.001 |
| *Response Mode* | 33.25 | <0.001 |
| *TE-Group\* Stimuli Condition* | 2.91 | 0.088 |
| *TE-Group\*Response Mode* | 0.01 | 0.925 |
| *Stimuli Condition\*Response Mode* | 14.7 | <0.001 |
| *TE-Group\* Stimuli Condition\*Response Mode* | 1.5 | 0.221 |

## References

1 Bitsko RH, Claussen AH, Lichstein J, Black LI, Jones SE, Danielson ML et al. Mental Health Surveillance Among Children — United States, 2013–2019. MMWR Suppl 2022; 71: 1–42.

2 Perou R, Bitsko RH, Blumberg SJ, Pastor P, Ghandour RM, Gfroerer JC et al. Mental health surveillance among children--United States, 2005-2011. MMWR Suppl 2013; 62: 1–35.

3 McLaughlin KA, Green JG, Gruber MJ, Sampson NA, Zaslavsky AM, Kessler RC. Childhood adversities and first onset of psychiatric disorders in a national sample of US adolescents. Arch Gen Psychiatry 2012; 69: 1151–1160.

4 Wehry AM, Beesdo-Baum K, Hennelly MM, Connolly SD, Strawn JR. Assessment and Treatment of Anxiety Disorders in Children and Adolescents. Curr Psychiat Rep 2015; 17: 52.

5 Teicher MH, Andersen SL, Polcari A, Anderson CM, Navalta CP, Kim DM. The neurobiological consequences of early stress and childhood maltreatment. Neurosci Biobehav Rev 2003; 27: 33–44.

6 Ellis BJ, Giudice MD. Beyond allostatic load: Rethinking the role of stress in regulating human development. Dev Psychopathol 2014; 26: 1–20.

7 Ellis BJ, Sheridan MA, Belsky J, McLaughlin KA. Why and how does early adversity influence development? Toward an integrated model of dimensions of environmental experience. Dev Psychopathol 2022; 34: 447–471.

8 Tottenham N, Hare TA, Quinn BT, Mccarry TW, Nurse M, Gilhooly T et al. Prolonged institutional rearing is associated with atypically large amygdala volume and difficulties in emotion regulation. Developmental Science 2010; 13: 46–61.

9 Lambert HK, King KM, Monahan KC, McLaughlin KA. Differential associations of threat and deprivation with emotion regulation and cognitive control in adolescence. Dev Psychopathol 2017; 29: 929–940.

10 Kim SG, Weissman DG, Sheridan MA, McLaughlin KA. Child abuse and automatic emotion regulation in children and adolescents. Dev Psychopathol 2023; 35: 157–167.

11 Hennessy KD, Rabideau GJ, Cicchetti D, Cummings EM. Responses of Physically Abused and Nonabused Children to Different Forms of Interadult Anger. Child Dev 1994; 65: 815–828.

12 Glaser J-P, Os J van, Portegijs PJM, Myin-Germeys I. Childhood trauma and emotional reactivity to daily life stress in adult frequent attenders of general practitioners. J Psychosom Res 2006; 61: 229–236.

13 McLaughlin KA, Kubzansky LD, Dunn EC, Waldinger R, Vaillant G, Koenen KC. Childhood Social Environment, Emotional Reactivity, to Stress, and Mood and Anxiety Disorders Across the Life Course. Depression and Anxiety 2010; 27: 1087–1094.

14 France JM, Reda M, Marusak HA, Riser M, Wiltshire CN, Davie WM et al. Anxiety, fear extinction, and threat-related amygdala reactivity in children exposed to urban trauma. J Exp Psychopathol 2022; 13: 20438087221132501.

15 Stevens JS, Rooij SJHV, Stenson AF, Ely TD, Powers A, Clifford A et al. Amygdala responses to threat in violence-exposed children depend on trauma context and maternal caregiving. Development and Psychopathology 2021;: 1–12.

16 Marusak HA, Zundel CG, Brown S, Rabinak CA, Thomason ME. Convergent behavioral and corticolimbic connectivity evidence of a negativity bias in children and adolescents. Soc Cogn Affect Neurosci 2017; 12: 517–525.

17 Hein TC, Goetschius LG, McLoyd VC, Brooks-Gunn J, McLanahan SS, Mitchell C et al. Childhood violence exposure and social deprivation are linked to adolescent threat and reward neural function. Social Cognitive and Affective Neuroscience 2020; 15: 1252–1259.

18 Marusak HA, Martin KR, Etkin A, Thomason ME. Childhood Trauma Exposure Disrupts the Automatic Regulation of Emotional Processing. Neuropsychopharmacology 2015; 40: 1250–1258.

19 McLaughlin KA, Peverill M, Gold AL, Alves S, Sheridan MA. Child Maltreatment and Neural Systems Underlying Emotion Regulation. J Am Acad Child Adolesc Psychiatry 2015; 54: 753–762.

20 Hare TA, Tottenham N, Galvan A, Voss HU, Glover GH, Casey BJ. Biological Substrates of Emotional Reactivity and Regulation in Adolescence During an Emotional Go-Nogo Task. Biological Psychiatry 2008; 63: 927–934.

21 Wessel JR, Anderson MC. Neural mechanisms of domain-general inhibitory control. Trends Cogn Sci 2024; 28: 124–143.

22 Bij J van der, Kelder RO den, Montagne B, Hagenaars MA. Inhibitory control in trauma-exposed youth: A systematic review. Neuroscience and Biobehavioral Reviews 2020; 118: 451–462.

23 Kelder RO den, Akker ALV den, Geurts HM, Lindauer RJL, Overbeek G. Executive functions in trauma-exposed youth: a meta-analysis. Eur J Psychotraumatology 2018; 9: 1450595.

24 Ordaz SJ, Foran W, Velanova K, Luna B. Longitudinal Growth Curves of Brain Function Underlying Inhibitory Control through Adolescence. J Neurosci 2013; 33: 18109–18124.

25 Tervo-Clemmens B, Calabro FJ, Parr AC, Fedor J, Foran W, Luna B. A canonical trajectory of executive function maturation from adolescence to adulthood. Nat Commun 2023; 14: 6922.

26 Quach A, Tervo-Clemmens B, Foran W, Calabro FJ, Chung T, Clark DB et al. Adolescent development of inhibitory control and substance use vulnerability: A longitudinal neuroimaging study. Dev Cogn Neurosci 2020; 42: 100771.

27 Tottenham N, Hare TA, Casey BJ. Behavioral assessment of emotion discrimination, emotion regulation, and cognitive control in childhood, adolescence, and adulthood. Frontiers in Psychology 2011; 2: 1–9.

28 Rocha GS, Freire MAM, Britto AM, Paiva KM, Oliveira RF, Fonseca IAT et al. Basal ganglia for beginners: the basic concepts you need to know and their role in movement control. Front Syst Neurosci 2023; 17: 1242929.

29 Paus T. Primate anterior cingulate cortex: Where motor control, drive and cognition interface. Nat Rev Neurosci 2001; 2: 417–424.

30 Rubia K, Smith AB, Woolley J, Nosarti C, Heyman I, Taylor E et al. Progressive increase of frontostriatal brain activation from childhood to adulthood during event-related tasks of cognitive control. Hum Brain Mapp 2006; 27: 973–993.

31 Rubia K, Smith AB, Taylor E, Brammer M. Linear age-correlated functional development of right inferior fronto-striato-cerebellar networks during response inhibition and anterior cingulate during error-related processes. Hum Brain Mapp 2007; 28: 1163–1177.

32 Asemi A, Ramaseshan K, Burgess A, Diwadkar VA, Bressler SL. Dorsal anterior cingulate cortex modulates supplementary motor area in coordinated unimanual motor behavior. Front Hum Neurosci 2015; 9: 309.

33 Williams ZM, Bush G, Rauch SL, Cosgrove GR, Eskandar EN. Human anterior cingulate neurons and the integration of monetary reward with motor responses. Nat Neurosci 2004; 7: 1370–1375.

34 Luppino G, Matelli M, Camarda RM, Gallese V, Rizzolatti G. Multiple representations of body movements in mesial area 6 and the adjacent cingulate cortex: An intracortical microstimulation study in the macaque monkey. J Comp Neurol 1991; 311: 463–482.

35 Teicher MH, Samson JA, Anderson CM, Ohashi K. The effects of childhood maltreatment on brain structure, function and connectivity. Nat Rev Neurosci 2016; 17: 652–666.

36 Koush Y, Rothman DL, Behar KL, Graaf RA de, Hyder F. Human brain functional MRS reveals interplay of metabolites implicated in neurotransmission and neuroenergetics. J Cereb Blood Flow Metab 2022; 42: 911–934.

37 Rothman DL, Graaf RA, Hyder F, Mason GF, Behar KL, Feyter HMD. In vivo 13C and 1H-[13C] MRS studies of neuroenergetics and neurotransmitter cycling, applications to neurological and psychiatric disease and brain cancer. NMR Biomed 2019; 32: e4172.

38 Mangia S, Tkáč I, Gruetter R, Moortele P-FVD, Giove F, Maraviglia B et al. Sensitivity of single-voxel 1H-MRS in investigating the metabolism of the activated human visual cortex at 7 T. Magn Reson Imaging 2006; 24: 343–348.

39 Hertz L, Xu J, Song D, Du T, Yan E, Peng L. Brain glycogenolysis, adrenoceptors, pyruvate carboxylase, Na+,K+-ATPase and Marie E. Gibbs’ pioneering learning studies. Front Integr Neurosci 2013; 7: 20.

40 Schousboe A, Bak LK, Waagepetersen HS. Astrocytic Control of Biosynthesis and Turnover of the Neurotransmitters Glutamate and GABA. Front Endocrinol 2013; 4: 102.

41 Tatti R, Haley MS, Swanson OK, Tselha T, Maffei A. Neurophysiology and Regulation of the Balance Between Excitation and Inhibition in Neocortical Circuits. Biol Psychiat 2017; 81: 821–831.

42 Zhou X, Zhu D, King SG, Lees CJ, Bennett AJ, Salinas E et al. Behavioral response inhibition and maturation of goal representation in prefrontal cortex after puberty. Proc Natl Acad Sci 2016; 113: 3353–3358.

43 Gonzalez-Burgos G, Miyamae T, Reddy N, Dawkins S, Chen C, Hill A et al. Mechanisms regulating the properties of inhibition-based gamma oscillations in primate prefrontal and parietal cortices. Cereb Cortex 2023; 33: 7754–7770.

44 Wang Z, Singh B, Zhou X, Constantinidis C. Strong Gamma Frequency Oscillations in the Adolescent Prefrontal Cortex. J Neurosci 2022; 42: 2917–2929.

45 Logothetis NK. What we can do and what we cannot do with fMRI. Nature 2008; 453: 869–878.

46 Stanley JA, Raz N. Functional Magnetic Resonance Spectroscopy: The “New” MRS for Cognitive Neuroscience and Psychiatry Research. Frontiers Psychiatry 2018; 9: 76.

47 Pasanta D, He JL, Ford T, Oeltzschner G, Lythgoe DJ, Puts NA. Functional MRS Studies of GABA and Glutamate/Glx – A Systematic Review and Meta-analysis. Neurosci Biobehav Rev 2022; 144: 104940.

48 Taylor R, Schaefer B, Densmore M, Neufeld RWJ, Rajakumar N, Williamson PC et al. Increased glutamate levels observed upon functional activation in the anterior cingulate cortex using the Stroop Task and functional spectroscopy. Neuroreport 2015; 26: 107–112.

49 Taylor R, Neufeld RWJ, Schaefer B, Densmore M, Rajakumar N, Osuch EA et al. Functional magnetic resonance spectroscopy of glutamate in schizophrenia and major depressive disorder: anterior cingulate activity during a color-word Stroop task. npj Schizophr 2015; 1: 15028.

50 France JM, Valbrun SA, Grasser LR, Wiltshire C, Basarkod S, Davie WM et al. Trauma from the Eye of the Beholder: Reporter Discordance in Child’s Trauma, Psychopathology, and Neurobiology. Biol Psychiatry: Cogn Neurosci Neuroimaging 2025. doi:10.1016/j.bpsc.2025.08.007.

51 Ford JD, R. Racusin, Rogers K, Ellis C, Schiffman J, Ribbe D et al. Traumatic events screening inventory for children (TESI-C) Version 8.4. National Center for PTSD and Dartmouth Child Psychiatry Research Group 2002.

52 Ghosh-Ippen C, Ford J, Racusin R, Acker M, Bosquet K, Rogers C et al. Traumatic Events Screening Inventory - Parent Report Revised. National Center for PTSD and Dartmouth Child Psychiatry Research Group 2002.

53 Choi KR, McCreary M, Ford JD, Koushkaki SR, Kenan KN, Zima BT. Validation of the Traumatic Events Screening Inventory for ACEs. Pediatrics 2019; 143: e20182546.

54 Copeland WE, Keeler G, Angold A, Costello EJ. Traumatic Events and Posttraumatic Stress in Childhood. Arch Gen Psychiatry 2007; 64: 577–584.

55 Kessler RC, McLaughlin KA, Green JG, Gruber MJ, Sampson NA, Zaslavsky AM et al. Childhood adversities and adult psychopathology in the WHO World Mental Health Surveys. Br J Psychiatry 2010; 197: 378–385.

56 Green JG, McLaughlin KA, Berglund PA, Gruber MJ, Sampson NA, Zaslavsky AM et al. Childhood Adversities and Adult Psychiatric Disorders in the National Comorbidity Survey Replication I: Associations With First Onset of DSM-IV Disorders. Arch Gen Psychiat 2010; 67: 113–123.

57 Copeland WE, Shanahan L, Costello EJ, Angold A. Childhood and Adolescent Psychiatric Disorders as Predictors of Young Adult Disorders. Arch Gen Psychiatry 2009; 66: 764–772.

58 Felitti VJ, Anda RF, Nordenberg D, Williamson DF, Spitz AM, Edwards V et al. Relationship of Childhood Abuse and Household Dysfunction to Many of the Leading Causes of Death in Adults The Adverse Childhood Experiences (ACE) Study. Am J Prev Med 1998; 14: 245–258.

59 Birmaher B, Brent DA, Chiappetta L, Bridge J, Monga S, Baugher M. Psychometric Properties of the Screen for Child Anxiety Related Emotional Disorders (SCARED): A Replication Study. J Am Acad Child Adolesc Psychiatry 1999; 36: 545–553.

60 Tottenham N, Tanaka JW, Leon AC, McCarry T, Nurse M, Hare TA et al. The NimStim set of facial expressions: Judgments from untrained research participants. Psychiatry Research 2009; 168: 242–249.

61 Lynn J, Woodcock EA, Anand C, Khatib D, Stanley JA. Differences in steady-state glutamate levels and variability between “non-task-active” conditions: Evidence from 1H fMRS of the prefrontal cortex. NeuroImage 2017; 172: 554–561.

62 Woodcock EA, Arshad M, Khatib D, Stanley JA. Automated Voxel Placement: A Linux-based Suite of Tools for Accurate and Reliable Single Voxel Coregistration. J Neuroimaging Psychiatry Neurology 2018; 3: 1–8.

63 Tkáć I, Gruetter R. Methodology of1H NMR spectroscopy of the human brain at very high magnetic fields. Appl Magn Reson 2004; 29: 139.

64 Gruetter R. Automatic, localized in Vivo adjustment of all first-and second-order shim coils. Magn Reson Med 1993; 29: 804–811.

65 Gasparovic C, Song T, Devier D, Bockholt HJ, Caprihan A, Mullins PG et al. Use of tissue water as a concentration reference for proton spectroscopic imaging. Magnet Reson Med 2006; 55: 1219–1226.

66 Provencher SW. Estimation of metabolite concentrations from localized in vivo proton NMR spectra. Magn Reson Med 1993; 30: 672–679.

67 Stanley JA, Drost DJ, Williamson PC, Thompson RT. The use of a priori knowledge to quantify short echo in vivo 1h mr spectra. Magn Reson Med 1995; 34: 17–24.

68 Govindaraju V, Young K, Maudsley AA. Proton NMR chemical shifts and coupling constants for brain metabolites. NMR in Biomedicine 2000; 13: 129–153.

69 Gasparovic C, Song T, Devier D, Bockholt HJ, Caprihan A, Mullins PG et al. Use of tissue water as a concentration reference for proton spectroscopic imaging. Magnet Reson Med 2006; 55: 1219–1226.

70 Posse S, Otazo R, Caprihan A, Bustillo J, Chen H, Henry P et al. Proton echo-planar spectroscopic imaging of J-coupled resonances in human brain at 3 and 4 Tesla. Magn Reson Med 2007; 58: 236–244.

71 Erdfelder E, Faul F, Buchner A. GPOWER: A general power analysis program. *Behav Res Methods, Instrum*, Comput 1996; 28: 1–11.

72 Etkin A, Egner T, Kalisch R. Emotional processing in anterior cingulate and medial prefrontal cortex. Trends Cogn Sci 2011; 15: 85–93.

73 Shin LM, Liberzon I. The Neurocircuitry of Fear, Stress, and Anxiety Disorders. Neuropsychopharmacol 2010; 35: 169–191.

74 Radley JJ, Rocher AB, Janssen WGM, Hof PR, McEwen BS, Morrison JH. Reversibility of apical dendritic retraction in the rat medial prefrontal cortex following repeated stress. Exp Neurol 2005; 196: 199–203.

75 Radley JJ, Rocher AB, Rodriguez A, Ehlenberger DB, Dammann M, McEwen BS et al. Repeated stress alters dendritic spine morphology in the rat medial prefrontal cortex. J Comp Neurol 2008; 507: 1141–1150.

76 Cerqueira JJ, Pêgo JM, Taipa R, Bessa JM, Almeida OFX, Sousa N. Morphological Correlates of Corticosteroid-Induced Changes in Prefrontal Cortex-Dependent Behaviors. J Neurosci 2005; 25: 7792–7800.

77 Goldwater DS, Pavlides C, Hunter RG, Bloss EB, Hof PR, McEwen BS et al. Structural and functional alterations to rat medial prefrontal cortex following chronic restraint stress and recovery. Neuroscience 2009; 164: 798–808.

78 Castillo-Gómez E, Pérez-Rando M, Bellés M, Gilabert-Juan J, Llorens JV, Carceller H et al. Early Social Isolation Stress and Perinatal NMDA Receptor Antagonist Treatment Induce Changes in the Structure and Neurochemistry of Inhibitory Neurons of the Adult Amygdala and Prefrontal Cortex. eNeuro 2017; 4: ENEURO.0034-17.2017.

79 Gildawie KR, Honeycutt JA, Brenhouse HC. Region-specific Effects of Maternal Separation on Perineuronal Net and Parvalbumin-expressing Interneuron Formation in Male and Female Rats. Neuroscience 2020; 428: 23–37.

80 Hawes MT, Szenczy AK, Klein DN, Hajcak G, Nelson BD. Increases in depression and anxiety symptoms in adolescents and young adults during the COVID-19 pandemic. Psychol Med 2022; 52: 3222–3230.

81 Merikangas KR, He JP, Burstein M, Swanson SA, Avenevoli S, Cui L et al. Lifetime prevalence of mental disorders in U.S. adolescents: Results from the national comorbidity survey replication-adolescent supplement (NCS-A). Journal of the American Academy of Child & Adolescent Psychiatry 2010; 49: 980–989.

82 Pollak SD, Sinha P. Effects of Early Experience on Children’s Recognition of Facial Displays of Emotion. Dev Psychol 2002; 38: 784–791.

83 Lambert HK, Sheridan MA, Sambrook KA, Rosen ML, Askren MK, McLaughlin KA. Hippocampal Contribution to Context Encoding across Development Is Disrupted following Early-Life Adversity. J Neurosci 2017; 37: 1925–1934.

84 Hertz L. The glutamate-glutamine (GABA) cycle: importance of late postnatal development and potential reciprocal interactions between biosynthesis and degradation. Frontiers in Endocrinology 2013. doi:10.3389/fendo.2013.00059.

